# Paucity of adaptive selection in PmrAB two-component system may resist emergence of colistin resistance in *Acinetobacter baumannii*

**DOI:** 10.1101/753665

**Authors:** Sudhakar Pagal, Rajagopalan Saranathan, Anshu Rani, Archana Tomar, K. P. Arunkumar, K. Prashanth

**Affiliations:** Nosocomial Pathogen Biology Lab, Department of Biotechnology, School of Life Sciences, Pondicherry University, Puducherry, India; Laboratory of Molecular Genetics, Centre for DNA Fingerprinting and Diagnostics (CDFD), Hyderabad, India

**Keywords:** Colistin resistance, PmrAB, Point mutation, Selection analysis, Purifying selection, *Acinetobacter baumannii*

## Abstract

Investigations on the selection pressure acting on point mutations in PmrAB two-component system may provide insights into the future of colistin therapy in *Acinetobacter baumannii*, since mutations in *pmrAB* are implicated in colistin resistance. We performed adaptive selection analysis of *pmrAB* and compared with the available data on colistin resistant strains. We analysed PmrAB sequences in 3113 draft genomes of *A. baumannii* obtained from RefSeq database. Adaptive selection analysis was performed by two widely used programs namely, HyPhy and PAML. In addition, to examine the reliability of the approach, the same analysis was performed on *gyrA* of *Escherichia coli* and *Salmonella enterica*, since adaptive mutations on *gyrA* confer quinolone resistance. Mutations that had caused colistin resistance were found to be neither adaptive nor polymorphic, rather they occur at sites that are either under neutral or purifying selection. Strong negative evolutionary selection pressure is also observed at sites throughout both PmrA and PmrB. Sites with high levels of polymorphisms in PmrAB were found to be under neutral selection. Notably, there was no sign of positive selection. Some of them are rather deleterious. These conditions might be maintaining the incidence of colistin resistance in *A. baumannii* under check. Therefore, in the context of colistin resistance, natural selection plays only a minor role and we assert that in future, *A. baumannii* may not be able to sustain and successfully disseminate colistin resistance. Therefore, at present the concerns raised about continuing the usage of colistin for the treatment against *A. baumannii* infections appears to be unnecessary.

## Introduction

Single nucleotide polymorphisms (SNPs) contribute greatly towards the microevolution of a bacterium (Chouard, 2010; Reznick & Ricklefs, 2009). Consequently, these bacteria are able to adapt to their environment comparatively faster without any significant associated costs. These mutations also have a profound effect on drug resistance and pathogenesis. In recent years, colistin resistance has been found to be due to SNPs in the genes encoding the two-component system (TCS) PmrAB, as well as in the *lpx* operon, that are responsible for the production and regulation of lipopolysaccharide (LPS) in bacterial cell walls (Moffatt et al., 2010; Snitkin et al., 2013; Thi Khanh Nhu et al., 2016).

PmrAB is thought to regulate lipid-A biosynthesis of gram negative outer membranes (Boll et al., 2016). Lipid-A is a constituent of the outer membrane LPS, which is the target of colistin. Colistin resistant strains of *A. baumannii* have modifications and/or loss of LPS from their outer membrane, with concomitant mutations in the *pmrCAB* and/or *lpx* operon (Boll et al., 2016; Carretero-Ledesma et al., 2018). Emergence of such resistance is worrisome, since colistin is the last-choice drug against multidrug resistant *A. baumannii*. The objective of this study was therefore to identify the hyper-variable regions of the TCS PmrAB in *A. baumannii* to gain insights into their contribution towards the structure and function of this TCS. Since point mutations in PmrAB have been implicated in resistance to colistin, sequence analysis of PmrAB in the context of the evolutionary selection pressure acting on these mutations may provide insights into the fate of colistin resistance in *A. baumannii*. We report that, both PmrA and PmrB are under strong purifying evolutionary selection pressure. In addition, there was no sign of adaptive evolution. Mutations that cause colistin resistance are neither adaptive nor polymorphic. Some of them are rather deleterious. Therefore, it is highly unlikely that these mutations will sustain in the long term and hence, *A. baumannii* may not be able to emerge as a colistin-resistant organism.

## Methods

Minimum inhibitory concentration (MIC) for colistin were performed according to European Committee on Antimicrobial Susceptibility Testing guidelines (EUCAST, 2017). *pmrA* coding sequences were amplified by *in house* designed primers and custom sequenced (**Supplementary methods**). For selection analysis, 3113 genomes were retrieved from RefSeq from which, *pmrA* and *pmrB* sequences were extracted. *pmrAB* sequences originated from colistin resistant strains that appeared in the literature were also included. Selection analysis was performed with Single Likelihood Ancestor Counting method of the HyPhy package, and CodeML of the PAML package. All methods in detail have been provided in **Supplementary methods**.

## Results

### Mutation hotspots in PmrAB are not responsible for colistin resistance

In our observations, two of our clinical isolates, PKAB15 and PKAB19 were identified to be colistin resistant (Table 1). We amplified and performed sequencing of the *pmrA* genes from these two isolates. One non-synonymous mutation in each of these isolates was identified in *pmrA*, when compared to the reference strain *A. baumannii* ATCC17978, both of which (D10N in PKAB15, and R212L in PKAB19) have never been reported hitherto. Both these mutations change charged amino acids with uncharged ones which, we believe might have significant alterations on the structure of the protein to bring about colistin resistance.

**Table 1.**
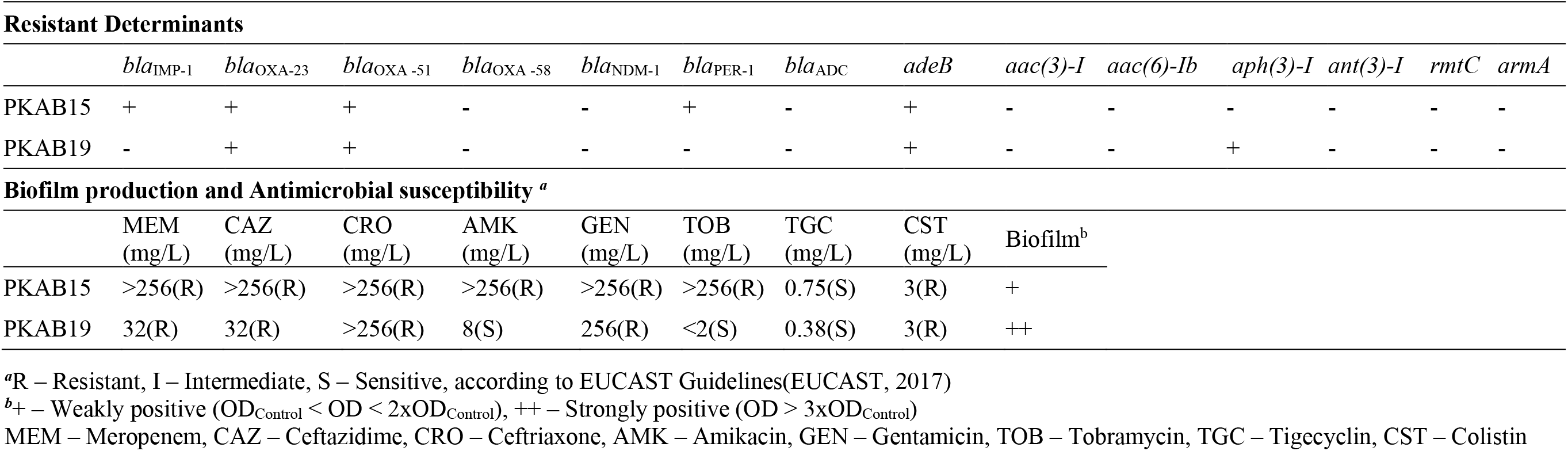
Molecular characters and other clinical details of isolates PKAB15 and PKAB19

Across all the 3113 genomes investigated in the study, 43 sites with non-synonymous mutations were identified in *pmrA* (Table 2). Of these, five were highly polymorphic sites. Most of the mutations occur in the signal receiver domain and the region connecting the receiver domain to the DNA-binding domain (Figure 1). The DNA binding domain recorded as few as seven sites with non-synonymous mutations, four of which had rare mutations while one was among the five most polymorphic sites. A well-conserved DNA binding domain emphasizes the importance of genes that are regulated by PmrAB, and therefore, it cannot afford many mutations in this region.

**Figure 1.**
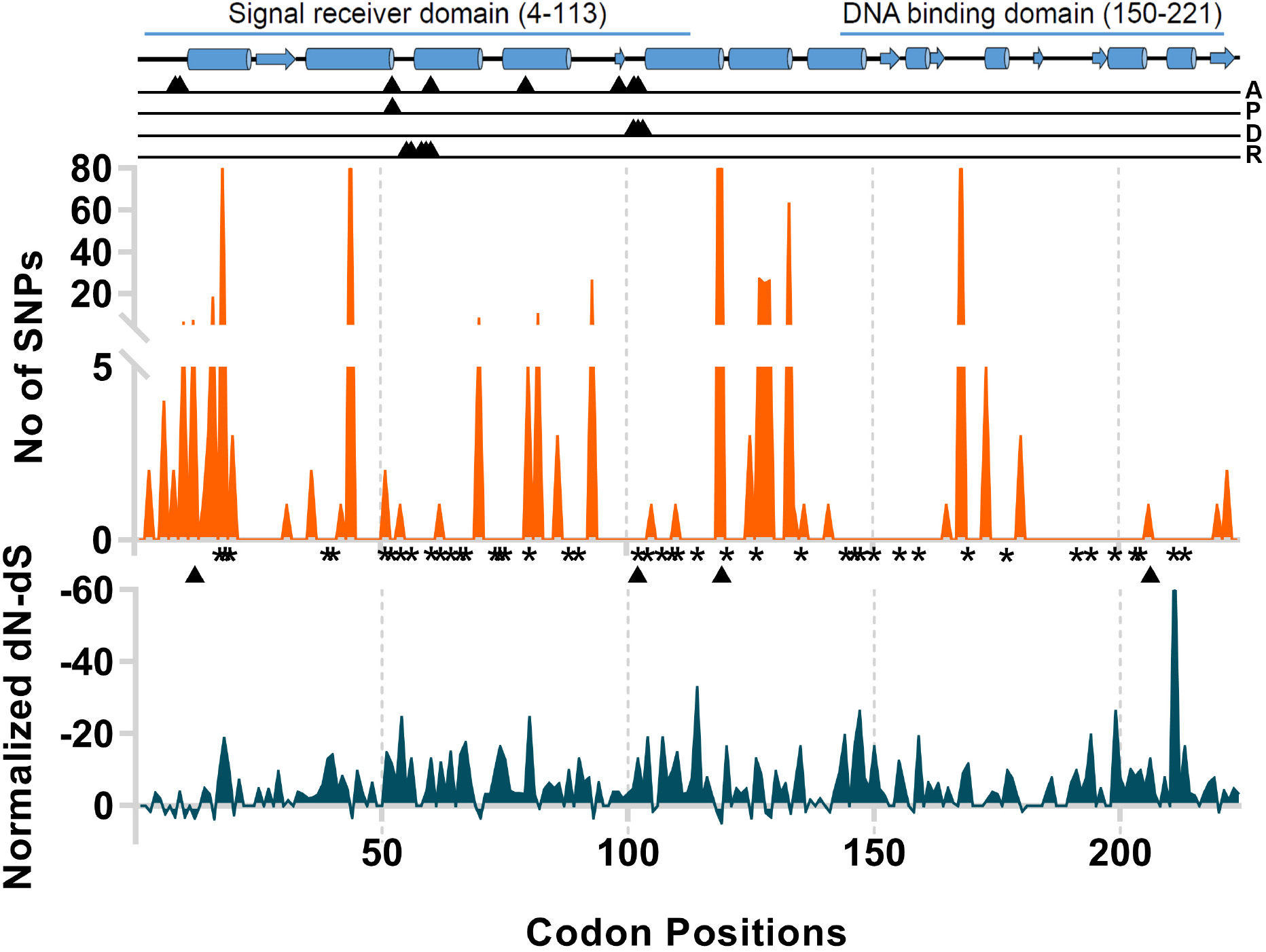
Mutation analysis of PmrA of *A. baumannii*. **Top panel** – The domain analysis (predicted by HMMER)(Finn et al., 2011). The secondary structure was predicted by Phyre2 web server (Kelly et al., 2015). The black triangles show functional sites of the protein (predicted by the Conserved Domain Database) (Marchler-Bauer et al., 2011). A-Active site, P-Phosphorylation site, D-Dimerization interface, R-Intermolecular recognition site. Barrels and arrows represent α-helices and β-sheets, respectively. Numbers in brackets denote the start-end positions of the corresponding domains. **Middle panel** – Positions and numbers of non-synonymous mutations across all the genomes under the analysis. **Bottom panel** – Selection along the protein sequence expressed in terms of normalized *dN – dS* (calculated by Datamonkey web server) (Kosakovsky Pond & Frost, 2005). Asterisks indicate sites under negative selection with p≤0·05. Triangles show the sites identified in colistin resistant isolates in previous literatures. This figure appears in colour in the online version of *JAC* and in black and white in the print version of *JAC*.

**Table 2.**
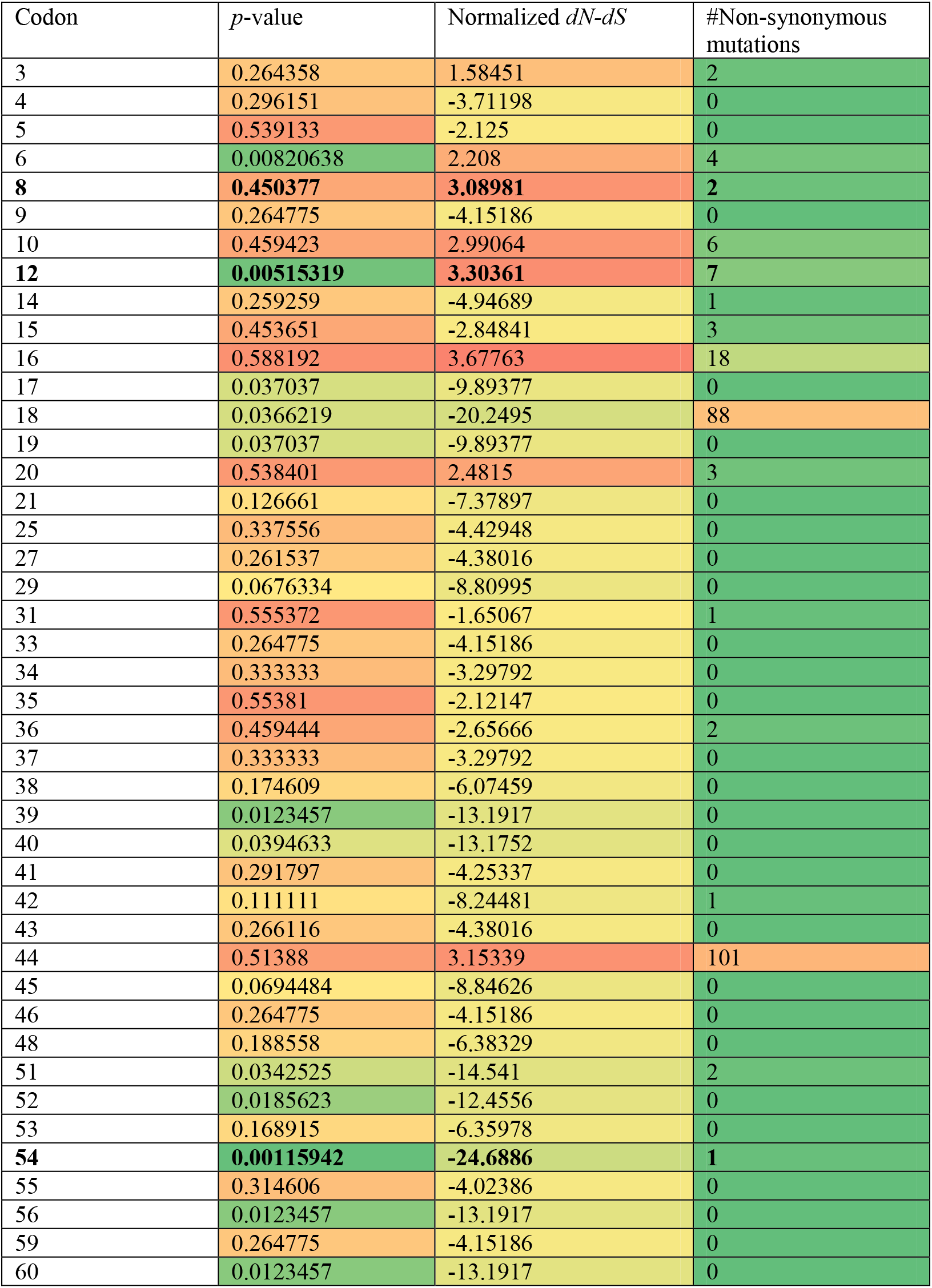
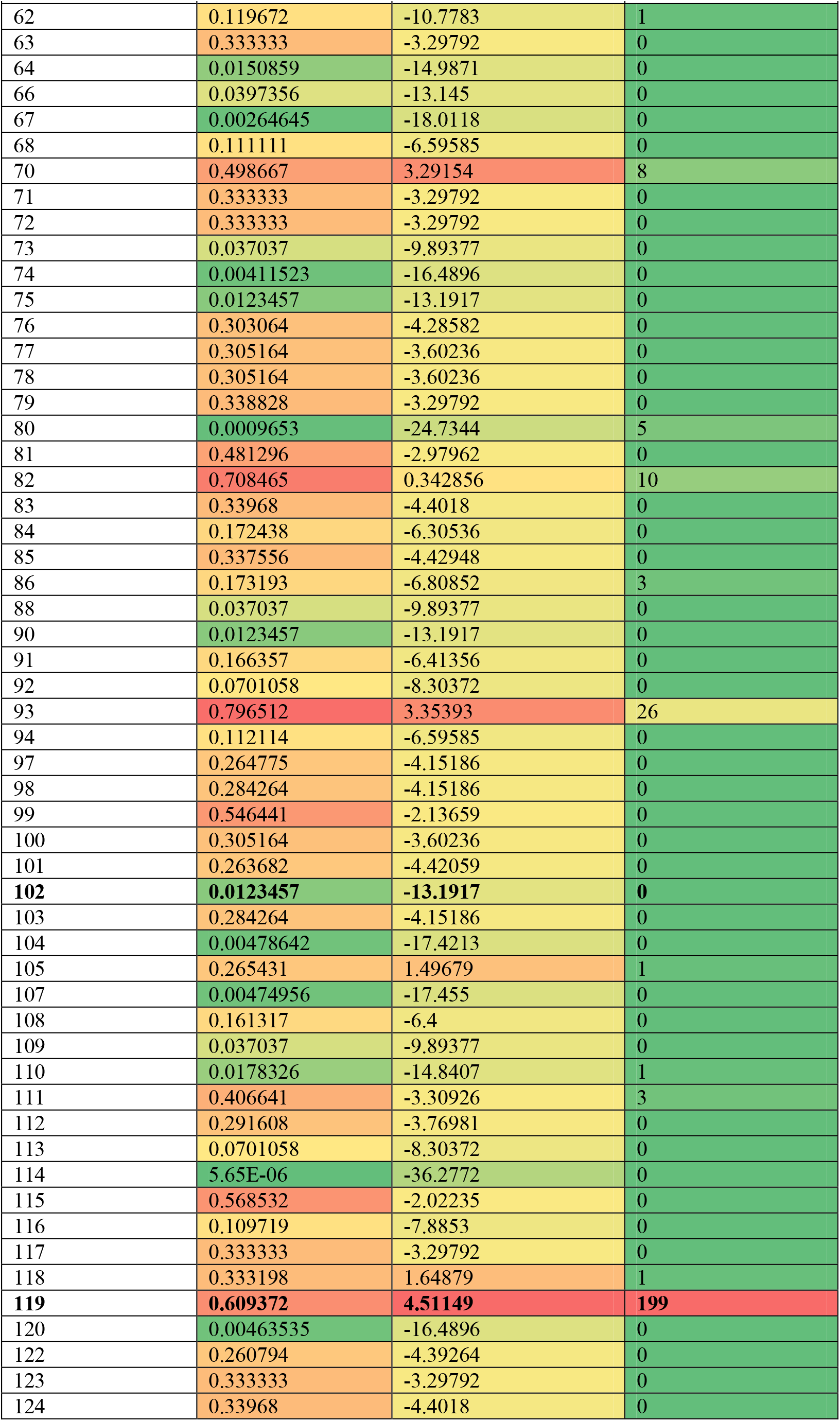
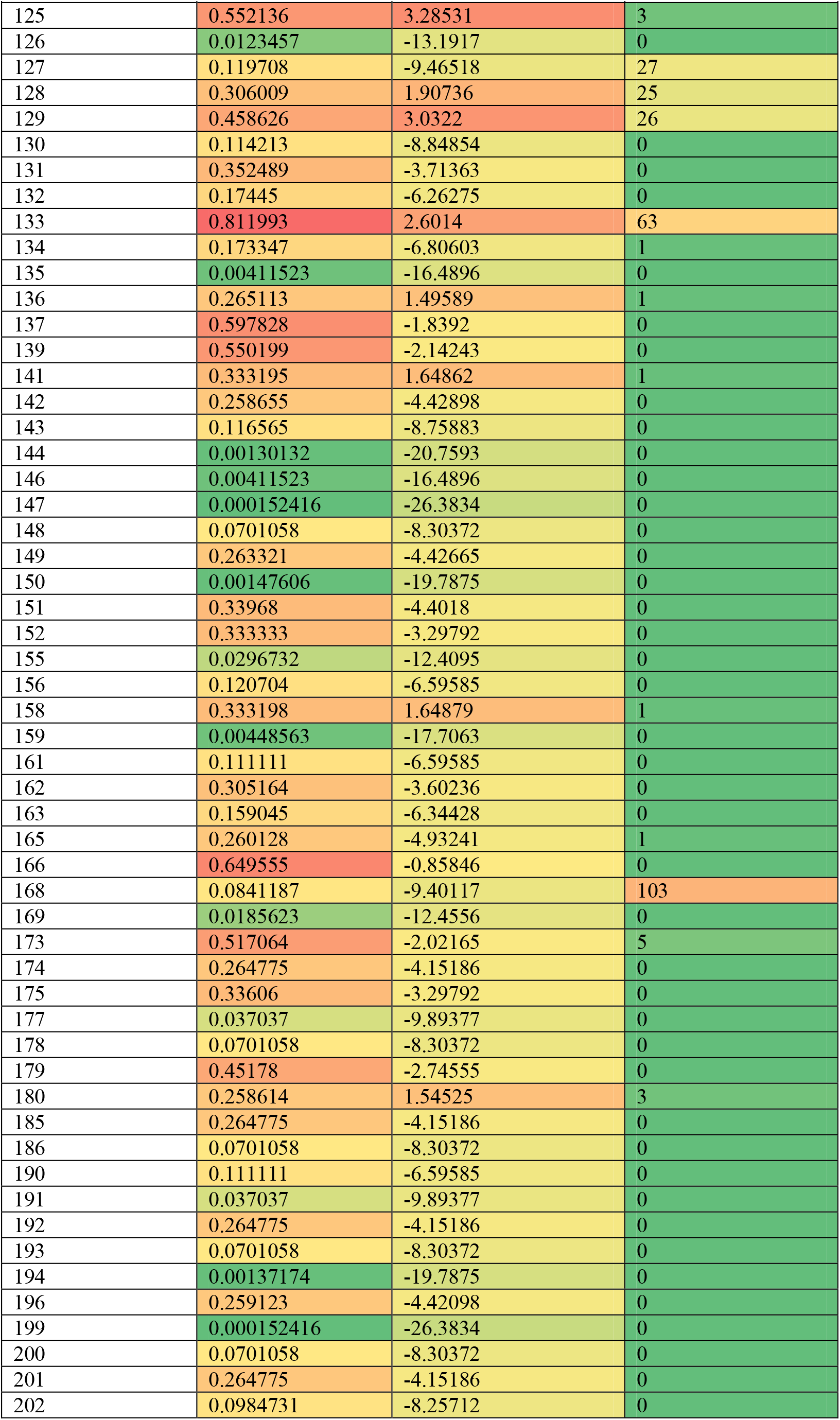
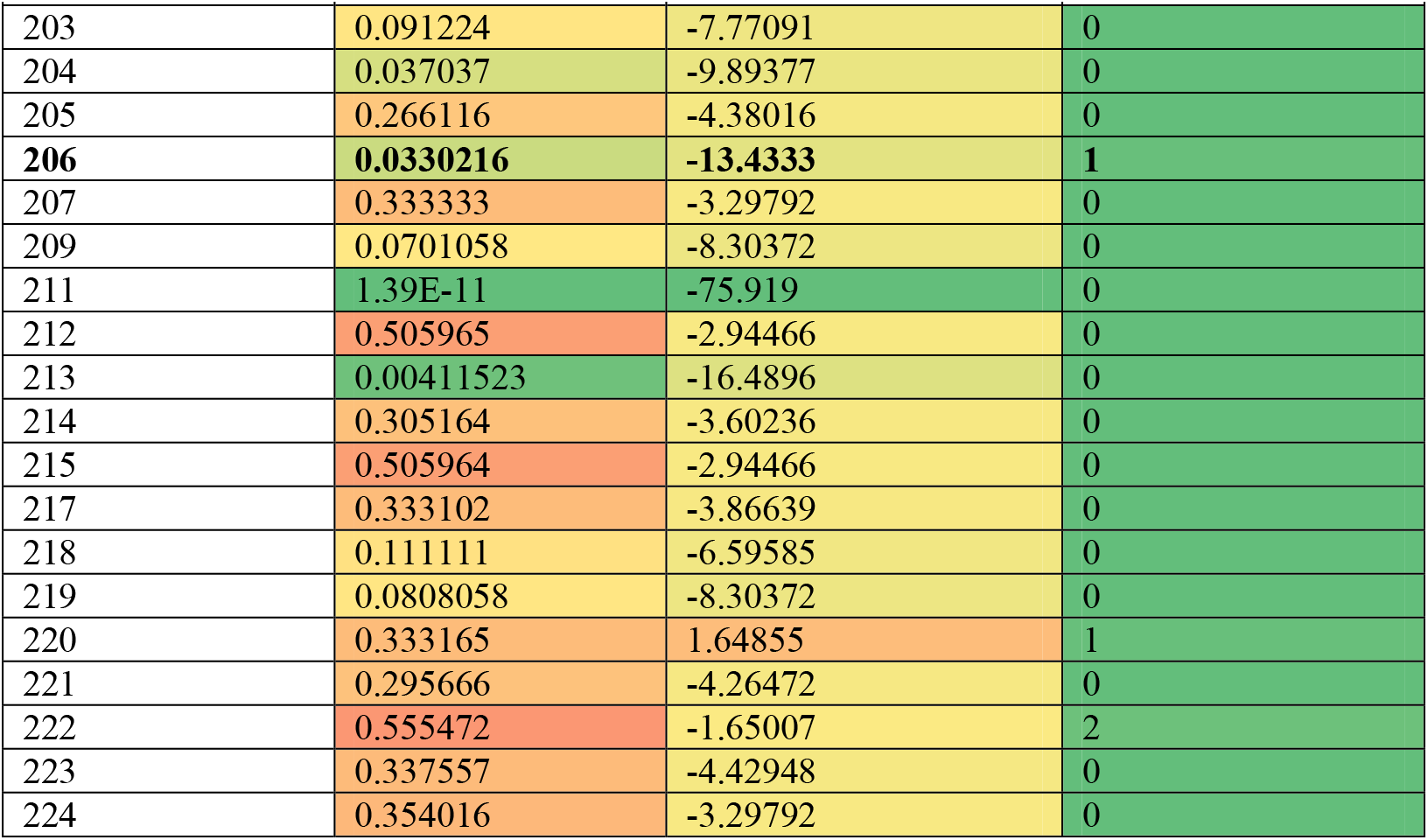
Codon-wise mutation and selection analysis and the corresponding *p*-values for *pmrA*. Only values for codons with non-zero normalized *dN* – *dS* values are shown. Sites reported to be associated with colistin resistant strains are shown in bold. *p*-values >0.05 are shaded red, <0.05 shaded green, intermediate color is yellow. Normalized *dN – d*S values >0 are shaded red, <0 shaded green, intermediate color is yellow.

| Codon | *p*-value | Normalized *dN-dS* | #Non-synonymous mutations |
| --- | --- | --- | --- |
| 3 | 0.264358 | 1.58451 | 2 |
| 4 | 0.296151 | -3.71198 | 0 |
| 5 | 0.539133 | -2.125 | 0 |
| 6 | 0.00820638 | 2.208 | 4 |
| **8** | **0.450377** | **3.08981** | **2** |
| 9 | 0.264775 | -4.15186 | 0 |
| 10 | 0.459423 | 2.99064 | 6 |
| **12** | **0.00515319** | **3.30361** | **7** |
| 14 | 0.259259 | -4.94689 | 1 |
| 15 | 0.453651 | -2.84841 | 3 |
| 16 | 0.588192 | 3.67763 | 18 |
| 17 | 0.037037 | -9.89377 | 0 |
| 18 | 0.0366219 | -20.2495 | 88 |
| 19 | 0.037037 | -9.89377 | 0 |
| 20 | 0.538401 | 2.4815 | 3 |
| 21 | 0.126661 | -7.37897 | 0 |
| 25 | 0.337556 | -4.42948 | 0 |
| 27 | 0.261537 | -4.38016 | 0 |
| 29 | 0.0676334 | -8.80995 | 0 |
| 31 | 0.555372 | -1.65067 | 1 |
| 33 | 0.264775 | -4.15186 | 0 |
| 34 | 0.333333 | -3.29792 | 0 |
| 35 | 0.55381 | -2.12147 | 0 |
| 36 | 0.459444 | -2.65666 | 2 |
| 37 | 0.333333 | -3.29792 | 0 |
| 38 | 0.174609 | -6.07459 | 0 |
| 39 | 0.0123457 | -13.1917 | 0 |
| 40 | 0.0394633 | -13.1752 | 0 |
| 41 | 0.291797 | -4.25337 | 0 |
| 42 | 0.111111 | -8.24481 | 1 |
| 43 | 0.266116 | -4.38016 | 0 |
| 44 | 0.51388 | 3.15339 | 101 |
| 45 | 0.0694484 | -8.84626 | 0 |
| 46 | 0.264775 | -4.15186 | 0 |
| 48 | 0.188558 | -6.38329 | 0 |
| 51 | 0.0342525 | -14.541 | 2 |
| 52 | 0.0185623 | -12.4556 | 0 |
| 53 | 0.168915 | -6.35978 | 0 |
| **54** | **0.00115942** | **-24.6886** | **1** |
| 55 | 0.314606 | -4.02386 | 0 |
| 56 | 0.0123457 | -13.1917 | 0 |
| 59 | 0.264775 | -4.15186 | 0 |
| 60 | 0.0123457 | -13.1917 | 0 |
| 62 | 0.119672 | -10.7783 | 1 |
| 63 | 0.333333 | -3.29792 | 0 |
| 64 | 0.0150859 | -14.9871 | 0 |
| 66 | 0.0397356 | -13.145 | 0 |
| 67 | 0.00264645 | -18.0118 | 0 |
| 68 | 0.111111 | -6.59585 | 0 |
| 70 | 0.498667 | 3.29154 | 8 |
| 71 | 0.333333 | -3.29792 | 0 |
| 72 | 0.333333 | -3.29792 | 0 |
| 73 | 0.037037 | -9.89377 | 0 |
| 74 | 0.00411523 | -16.4896 | 0 |
| 75 | 0.0123457 | -13.1917 | 0 |
| 76 | 0.303064 | -4.28582 | 0 |
| 77 | 0.305164 | -3.60236 | 0 |
| 78 | 0.305164 | -3.60236 | 0 |
| 79 | 0.338828 | -3.29792 | 0 |
| 80 | 0.0009653 | -24.7344 | 5 |
| 81 | 0.481296 | -2.97962 | 0 |
| 82 | 0.708465 | 0.342856 | 10 |
| 83 | 0.33968 | -4.4018 | 0 |
| 84 | 0.172438 | -6.30536 | 0 |
| 85 | 0.337556 | -4.42948 | 0 |
| 86 | 0.173193 | -6.80852 | 3 |
| 88 | 0.037037 | -9.89377 | 0 |
| 90 | 0.0123457 | -13.1917 | 0 |
| 91 | 0.166357 | -6.41356 | 0 |
| 92 | 0.0701058 | -8.30372 | 0 |
| 93 | 0.796512 | 3.35393 | 26 |
| 94 | 0.112114 | -6.59585 | 0 |
| 97 | 0.264775 | -4.15186 | 0 |
| 98 | 0.284264 | -4.15186 | 0 |
| 99 | 0.546441 | -2.13659 | 0 |
| 100 | 0.305164 | -3.60236 | 0 |
| 101 | 0.263682 | -4.42059 | 0 |
| **102** | **0.0123457** | **-13.1917** | **0** |
| 103 | 0.284264 | -4.15186 | 0 |
| 104 | 0.00478642 | -17.4213 | 0 |
| 105 | 0.265431 | 1.49679 | 1 |
| 107 | 0.00474956 | -17.455 | 0 |
| 108 | 0.161317 | -6.4 | 0 |
| 109 | 0.037037 | -9.89377 | 0 |
| 110 | 0.0178326 | -14.8407 | 1 |
| 111 | 0.406641 | -3.30926 | 3 |
| 112 | 0.291608 | -3.76981 | 0 |
| 113 | 0.0701058 | -8.30372 | 0 |
| 114 | 5.65E-06 | -36.2772 | 0 |
| 115 | 0.568532 | -2.02235 | 0 |
| 116 | 0.109719 | -7.8853 | 0 |
| 117 | 0.333333 | -3.29792 | 0 |
| 118 | 0.333198 | 1.64879 | 1 |
| **119** | **0.609372** | **4.51149** | **199** |
| 120 | 0.00463535 | -16.4896 | 0 |
| 122 | 0.260794 | -4.39264 | 0 |
| 123 | 0.333333 | -3.29792 | 0 |
| 124 | 0.33968 | -4.4018 | 0 |
| 125 | 0.552136 | 3.28531 | 3 |
| 126 | 0.0123457 | -13.1917 | 0 |
| 127 | 0.119708 | -9.46518 | 27 |
| 128 | 0.306009 | 1.90736 | 25 |
| 129 | 0.458626 | 3.0322 | 26 |
| 130 | 0.114213 | -8.84854 | 0 |
| 131 | 0.352489 | -3.71363 | 0 |
| 132 | 0.17445 | -6.26275 | 0 |
| 133 | 0.811993 | 2.6014 | 63 |
| 134 | 0.173347 | -6.80603 | 1 |
| 135 | 0.00411523 | -16.4896 | 0 |
| 136 | 0.265113 | 1.49589 | 1 |
| 137 | 0.597828 | -1.8392 | 0 |
| 139 | 0.550199 | -2.14243 | 0 |
| 141 | 0.333195 | 1.64862 | 1 |
| 142 | 0.258655 | -4.42898 | 0 |
| 143 | 0.116565 | -8.75883 | 0 |
| 144 | 0.00130132 | -20.7593 | 0 |
| 146 | 0.00411523 | -16.4896 | 0 |
| 147 | 0.000152416 | -26.3834 | 0 |
| 148 | 0.0701058 | -8.30372 | 0 |
| 149 | 0.263321 | -4.42665 | 0 |
| 150 | 0.00147606 | -19.7875 | 0 |
| 151 | 0.33968 | -4.4018 | 0 |
| 152 | 0.333333 | -3.29792 | 0 |
| 155 | 0.0296732 | -12.4095 | 0 |
| 156 | 0.120704 | -6.59585 | 0 |
| 158 | 0.333198 | 1.64879 | 1 |
| 159 | 0.00448563 | -17.7063 | 0 |
| 161 | 0.111111 | -6.59585 | 0 |
| 162 | 0.305164 | -3.60236 | 0 |
| 163 | 0.159045 | -6.34428 | 0 |
| 165 | 0.260128 | -4.93241 | 1 |
| 166 | 0.649555 | -0.85846 | 0 |
| 168 | 0.0841187 | -9.40117 | 103 |
| 169 | 0.0185623 | -12.4556 | 0 |
| 173 | 0.517064 | -2.02165 | 5 |
| 174 | 0.264775 | -4.15186 | 0 |
| 175 | 0.33606 | -3.29792 | 0 |
| 177 | 0.037037 | -9.89377 | 0 |
| 178 | 0.0701058 | -8.30372 | 0 |
| 179 | 0.45178 | -2.74555 | 0 |
| 180 | 0.258614 | 1.54525 | 3 |
| 185 | 0.264775 | -4.15186 | 0 |
| 186 | 0.0701058 | -8.30372 | 0 |
| 190 | 0.111111 | -6.59585 | 0 |
| 191 | 0.037037 | -9.89377 | 0 |
| 192 | 0.264775 | -4.15186 | 0 |
| 193 | 0.0701058 | -8.30372 | 0 |
| 194 | 0.00137174 | -19.7875 | 0 |
| 196 | 0.259123 | -4.42098 | 0 |
| 199 | 0.000152416 | -26.3834 | 0 |
| 200 | 0.0701058 | -8.30372 | 0 |
| 201 | 0.264775 | -4.15186 | 0 |
| 202 | 0.0984731 | -8.25712 | 0 |
| 203 | 0.091224 | -7.77091 | 0 |
| 204 | 0.037037 | -9.89377 | 0 |
| 205 | 0.266116 | -4.38016 | 0 |
| **206** | **0.0330216** | **-13.4333** | **1** |
| 207 | 0.333333 | -3.29792 | 0 |
| 209 | 0.0701058 | -8.30372 | 0 |
| 211 | 1.39E-11 | -75.919 | 0 |
| 212 | 0.505965 | -2.94466 | 0 |
| 213 | 0.00411523 | -16.4896 | 0 |
| 214 | 0.305164 | -3.60236 | 0 |
| 215 | 0.505964 | -2.94466 | 0 |
| 217 | 0.333102 | -3.86639 | 0 |
| 218 | 0.111111 | -6.59585 | 0 |
| 219 | 0.0808058 | -8.30372 | 0 |
| 220 | 0.333165 | 1.64855 | 1 |
| 221 | 0.295666 | -4.26472 | 0 |
| 222 | 0.555472 | -1.65007 | 2 |
| 223 | 0.337557 | -4.42948 | 0 |
| 224 | 0.354016 | -3.29792 | 0 |

One hundred fifty sites with amino acid substitutions in the PmrB protein sequence were found in our study, of which, 64 sites had rare mutations, each occurring in ≤0.1% of the genomes analysed (Table 3). Eighteen of the sites were highly polymorphic (Table 3 **&** Figure 2**, middle panel**). Most of these belong to the two trans-membrane regions and the intracellular domain connecting them. The HATPase_c domain, which binds the ATP moiety required for auto-phosphorylation of the sensor kinase, is the most conserved domain. In particular, with the exception of a few rare mutations, the ATP binding domain, Mg^2+^ binding domain and the G-X-G motif were well conserved (Figure 2**, upper & middle panel**). The Mg^2+^ metal cofactor and the G-X-G motif line the ATP binding pocket and are therefore crucial for the function of PmrB. Clearly, mutations in these sites can severely compromise its function.

**Figure 2.**
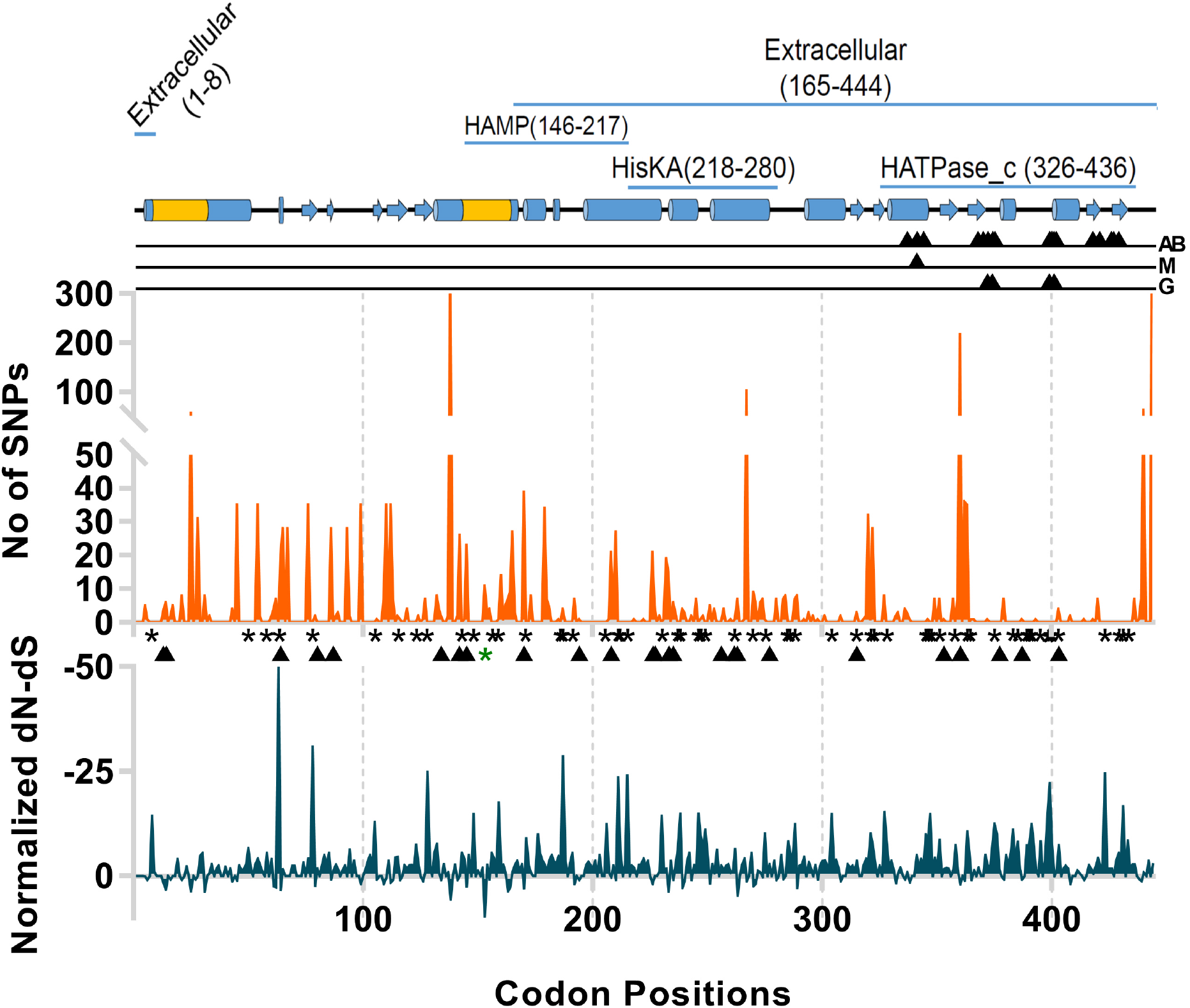
Mutation analysis of PmrB of *A. baumannii*. **Top panel** – The domain analysis (predicted by HMMER)(Finn et al., 2011). The secondary structure was predicted by Phyre2 server and trans-membrane (yellow barrels) by TMHMM (Kelly et al., 2015; Sonnhammer et al., 1998). The black triangles show functional sites of the protein (predicted by the Conserved Domain Database) (Marchler-Bauer et al., 2011). AB-ATP binding site, M-Mg^2+^ binding site, G-GXG motif. Barrels and arrows represent α-helices and β-sheets, respectively. Yellow coloured regions of the α-helices denote transmembrane domains. Numbers in brackets denote the start-end positions of the corresponding domains. **Middle panel** – Positions and numbers of non-synonymous mutations across all the genomes under the analysis. **Bottom panel** – Selection along the protein sequence in terms of normalized *dN – dS* (calculated by Datamonkey web server) (Kosakovsky Pond & Frost, 2005). Asterisks indicate sites under negative selection with p≤0·05, green asterisk representing a site with positive selection. Triangles show the sites identified in colistin resistant isolates in previous literatures. This figure appears in colour in the online version of *JAC* and in black and white in the print version of *JAC*.

**Table 3.** Codon-wise mutation and selection analysis and the corresponding *p*-values for *pmrB*. Only values for codons with non-zero normalized *dN* – *dS* values are shown. Sites reported to be associated with colistin resistant strains are shown in bold. *p*-values >0.05 are shaded red, <0.05 shaded green, intermediate color is yellow. Normalized *dN – d*S values >0 are shaded red, <0 shaded green, intermediate color is yellow.

| Codon | *p*-value | Normalized<br>*dN*-*dS* | #Non-synonymous<br>mutations |
| --- | --- | --- | --- |
| 5 | 0.75505 | 0.02736 | 5 |
| 6 | 0.23262 | 0.86962 | 1 |
| 8 | 0.00182 | -14.404 | 0 |
| 9 | 0.58075 | -1.0608 | 0 |
| 13 | **0.55552** | **1.84816** | **4** |
| 14 | **0.74388** | **3.18365** | **6** |
| 16 | 0.24758 | 0.81882 | 2 |
| 17 | **0.67942** | **-0.0326** | **5** |
| 19 | 0.08941 | -4.1207 | 0 |
| 21 | 0.33333 | 0.92411 | 8 |
| 23 | 0.27049 | -2.2776 | 0 |
| 25 | 0.75816 | 3.51406 | 57 |
| 27 | 0.25223 | 0.82388 | 3 |
| 29 | 0.07326 | -4.9802 | 0 |
| 30 | 0.05913 | -5.4891 | 0 |
| 31 | 0.80176 | 3.69314 | 8 |
| 33 | 0.58791 | -0.7609 | 2 |
| 34 | 0.28581 | -2.8642 | 0 |
| 36 | 0.33333 | -1.8482 | 0 |
| 37 | 0.24741 | -2.4901 | 0 |
| 40 | 0.3244 | -2.1937 | 0 |
| 42 | 0.33333 | -1.8482 | 0 |
| 43 | 0.28627 | 1.14615 | 4 |
| 45 | 0.59629 | 0.97052 | 15 |
| 46 | 0.27067 | -2.4901 | 0 |
| 47 | 0.5053 | -1.331 | 0 |
| 48 | 0.33333 | -1.8482 | 0 |
| 49 | 0.22739 | -2.8632 | 0 |
| 50 | 0.02931 | -6.6678 | 0 |
| 51 | 0.33333 | -1.8482 | 0 |
| 53 | 0.23685 | -2.7488 | 0 |
| 54 | 0.15783 | -4.1439 | 35 |
| 55 | 0.25926 | -2.7723 | 2 |
| 56 | 0.5046 | -1.3278 | 0 |
| 58 | 0.03704 | -5.5447 | 0 |
| 60 | 0.236 | -4.0995 | 2 |
| 61 | 0.57394 | 2.45633 | 4 |
| 62 | 0.56844 | 2.64003 | 7 |
| 63 | 7.39E-13 | -49.802 | 0 |
| 64 | **0.67964** | **3.27573** | **20** |
| 65 | 0.44231 | -1.6114 | 28 |
| 67 | 0.41978 | -0.0866 | 28 |
| 69 | 0.27067 | -2.4901 | 0 |

| Codon | *p*-value | Normalized<br>*dN-dS* | #Non-synonymous<br>mutations |
| --- | --- | --- | --- |
| 70 | 0.24741 | -2.4901 | 0 |
| 74 | 0.19416 | -3.1985 | 0 |
| 76 | 0.1748 | -4.2132 | 35 |
| 78 | 1.37E-08 | -30.905 | 0 |
| 79 | 0.0036 | 2.24631 | 2 |
| **80** | **0.33333** | **-1.8482** | **0** |
| 82 | 0.24247 | -2.7524 | 0 |
| 83 | 0.06121 | -4.9802 | 0 |
| 84 | 0.29742 | -2.7524 | 0 |
| 85 | 0.03704 | -5.5447 | 0 |
| 86 | 0.24741 | 0.81861 | 28 |
| **87** | **0.50023** | **-1.3073** | **0** |
| 88 | 0.26749 | -2.7265 | 2 |
| 89 | 0.43388 | -1.6695 | 3 |
| 91 | 0.45724 | -1.5569 | 0 |
| 92 | 0.24741 | -2.4901 | 0 |
| 93 | 0.74348 | -0.0338 | 28 |
| 94 | 0.07402 | -6.2937 | 2 |
| 96 | 0.11111 | -3.6965 | 0 |
| 99 | 0.82493 | 1.76685 | 35 |
| 101 | 0.33333 | -1.8482 | 0 |
| 102 | 0.12153 | -3.6965 | 0 |
| 103 | 0.06121 | -4.9802 | 0 |
| 104 | 0.27067 | -2.4901 | 0 |
| 105 | 0.00051 | -12.938 | 0 |
| 106 | 0.24743 | 0.81867 | 1 |
| 109 | 0.5373 | 1.81139 | 7 |
| 110 | 0.72533 | 0.40327 | 35 |
| 111 | 0.11122 | -3.6965 | 0 |
| 112 | 0.3285 | 1.52432 | 35 |
| 113 | 0.32791 | 0.91666 | 7 |
| 114 | 0.08543 | -4.5292 | 0 |
| 115 | 0.11109 | -4.621 | 2 |
| **116** | **0.27065** | **0.92403** | **1** |
| 118 | 0.25475 | -2.6568 | 0 |
| 119 | 0.40741 | -1.8482 | 4 |
| 120 | 0.25007 | -2.6131 | 0 |
| **121** | **0.24438** | **-2.7308** | **0** |
| 122 | 0.24741 | -2.4901 | 0 |
| 123 | 0.01983 | -7.4702 | 0 |
| 124 | 0.34697 | -3.1326 | 2 |
| 126 | 0.30008 | 1.16939 | 1 |
| 127 | 0.5483 | -0.9621 | 7 |
| 128 | 8.59E-07 | -24.901 | 0 |
| 129 | 0.29979 | -2.7306 | 0 |
| 131 | 0.05609 | -5.4979 | 0 |
| 132 | 0.04702 | -7.673 | 8 |
| 133 | 0.55556 | -0.9241 | 4 |
| **134** | **0.33321** | **0.92395** | **2** |
| 135 | 0.22887 | -2.8447 | 0 |
| 137 | 0.5741 | -0.8505 | 2 |
| 138 | 0.97399 | 5.54467 | 375 |
| 140 | 0.23685 | -2.7488 | 0 |
| 141 | 0.33331 | 0.92408 | 2 |
| 143 | 0.03704 | -5.5447 | 0 |
| 144 | 0.15348 | -4.159 | 1 |
| **145** | **0.00882** | **2.47068** | **23** |
| 146 | 0.06121 | -4.9802 | 0 |
| 148 | 0.00015 | -14.786 | 0 |
| 150 | 0.5046 | -1.3278 | 0 |
| 152 | 0.29902 | -2.0603 | 0 |
| 153 | 0.01367 | 9.48851 | 11 |
| 154 | 0.25926 | -2.7723 | 3 |
| 155 | 0.24759 | 0.8188 | 4 |
| 156 | 0.03704 | -5.5447 | 0 |
| 157 | 0.14703 | -4.19 | 0 |
| 159 | 0.00013 | -17.562 | 1 |
| 160 | 0.50737 | -1.3069 | 14 |
| 161 | 0.25926 | -2.7723 | 1 |
| 162 | 0.55556 | -0.9241 | 5 |
| 163 | 0.75834 | 3.51471 | 7 |
| 164 | 0.65538 | 2.63617 | 9 |
| 165 | 0.5529 | -0.9379 | 27 |
| 166 | 0.27169 | -2.6968 | 3 |
| 167 | 0.11111 | -3.6965 | 0 |
| 168 | 0.51381 | -1.2995 | 0 |
| 169 | 0.2346 | -2.8447 | 0 |
| **170** | **0.91393** | **2.80089** | **39** |
| 171 | 0.01864 | -9.0238 | 0 |
| 172 | 0.24741 | -2.4901 | 0 |
| 173 | 0.57387 | -0.852 | 8 |
| 176 | 0.00375 | -9.9603 | 0 |
| 177 | 0.05605 | -5.5 | 0 |
| 178 | 0.13718 | -3.8698 | 0 |
| 179 | 0.35446 | -2.7808 | 34 |
| 180 | 0.54467 | -1.1199 | 7 |
| 181 | 0.21056 | -3.6934 | 5 |
| 182 | 0.06121 | -4.9802 | 0 |
| **183** | **0.11419** | **-3.6965** | **0** |
| **184** | **0.22729** | **-2.8644** | **0** |
| 185 | 0.13346 | -4.8381 | 1 |
| 186 | 0.14473 | -4.0495 | 0 |
| 187 | 2.85E-06 | -28.648 | 4 |
| 188 | 0.11111 | -3.6965 | 0 |
| **190** | **0.22729** | **-2.8644** | **0** |
| 191 | 0.03704 | -5.5447 | 0 |
| **192** | **0.78192** | **1.79977** | **7** |
| **194** | **0.27072** | **0.92411** | **1** |
| 197 | 0.23685 | -2.7488 | 0 |
| 198 | 0.13218 | -3.4066 | 0 |
| 199 | 0.2602 | -2.6265 | 0 |
| 200 | 0.33333 | -1.8482 | 0 |
| 201 | 0.11111 | -3.6965 | 0 |
| 203 | 0.06121 | -4.9802 | 0 |
| 206 | 0.00093 | -12.45 | 0 |
| 207 | 0.5736 | -0.853 | 1 |
| **208** | **0.67468** | **0.21796** | **21** |
| 209 | 0.24741 | -2.4901 | 0 |
| 210 | 0.23891 | 0.8543 | 27 |
| 211 | 1.29E-06 | -23.529 | 0 |
| 212 | 0.02674 | -6.181 | 0 |
| 214 | 0.23391 | -2.8531 | 0 |
| 215 | 6.27E-07 | -24.027 | 0 |
| 217 | 0.24741 | -2.4901 | 0 |
| 218 | 0.40536 | -2.0004 | 1 |
| 220 | 0.24279 | -2.7488 | 0 |
| 221 | 0.28777 | -2.8447 | 0 |
| 222 | 0.15337 | -4.1615 | 1 |
| 223 | 0.29864 | -2.0629 | 0 |
| 224 | 0.11111 | -3.6965 | 0 |
| **226** | **0.86831** | **1.84822** | **21** |
| 229 | 0.53891 | -1.1768 | 2 |
| 230 | 0.00046 | -14.352 | 0 |
| 231 | 0.483 | -1.7181 | 1 |
| **232** | **0.42867** | **-1.8548** | **19** |
| **233** | **0.80247** | **3.69645** | **15** |
| **234** | **0.11111** | **-3.6965** | **0** |
| **235** | **0.80225** | **0.92229** | **6** |
| 236 | 0.25926 | -2.7723 | 3 |
| 237 | 0.03526 | -6.2927 | 0 |
| 238 | 0.00023 | -14.941 | 0 |
| 239 | 0.75517 | 2.72389 | 4 |
| 240 | 0.2854 | 1.14537 | 3 |
| 241 | 0.11111 | -3.6965 | 0 |
| 242 | 0.05965 | -5.465 | 0 |
| 243 | 0.29898 | 0.87883 | 2 |
| 244 | 0.25012 | -2.6126 | 0 |
| 245 | 0.55144 | -0.985 | 7 |
| 246 | 0.00023 | -14.941 | 0 |
| 247 | 0.03024 | -9.7957 | 1 |
| 248 | 0.05025 | -6.6009 | 2 |
| 249 | 0.00137 | -11.089 | 0 |
| 250 | 0.05688 | -5.4596 | 0 |
| 251 | 0.44218 | 1.7085 | 5 |
| 252 | 0.22712 | -2.8665 | 0 |
| 253 | 0.59374 | -0.7965 | 4 |
| 254 | 0.03541 | -6.2853 | 0 |
| **256** | **0.24741** | **-2.4901** | **0** |
| 257 | 0.50503 | -1.3142 | 4 |
| 258 | 0.24741 | -2.4901 | 0 |
| 259 | 0.2374 | -2.8111 | 0 |
| 260 | 0.55929 | -0.9126 | 1 |
| 261 | 0.50023 | -1.3073 | 0 |
| **262** | **0.04518** | **-6.4739** | **1** |
| **263** | **0.86792** | **4.61876** | **7** |
| 265 | 0.30006 | -2.7282 | 0 |
| 266 | 0.15337 | -4.1615 | 1 |
| 267 | 0.80615 | 1.23405 | 102 |
| 269 | 0.03704 | -5.5447 | 0 |
| 270 | 0.51364 | 2.34224 | 9 |
| 271 | 0.55531 | 1.84772 | 6 |
| **272** | **0.50568** | **-1.3328** | **0** |
| 273 | 0.5391 | -0.9241 | 6 |
| 274 | 0.67737 | 0.17619 | 7 |
| 275 | 0.00686 | -10.165 | 1 |
| 277 | 0.28777 | -2.8447 | 0 |
| 279 | 0.11111 | -3.6965 | 0 |
| 280 | 0.33333 | 0.92411 | 7 |
| 282 | 0.06121 | -4.9802 | 0 |
| 283 | 0.05268 | 0.65033 | 7 |
| 284 | 0.69625 | -0.0465 | 8 |
| 285 | 0.01324 | -8.2554 | 0 |
| 286 | 0.03704 | -5.5447 | 0 |
| 287 | 0.11111 | -3.6965 | 0 |
| 288 | 0.0025 | -12.454 | 7 |
| 289 | 0.6288 | 2.772 | 7 |
| 290 | 0.24741 | -2.4901 | 0 |
| 291 | 0.28816 | -2.8408 | 0 |
| 292 | 0.11121 | -4.6185 | 1 |
| 294 | 0.25765 | -3.3392 | 3 |
| 295 | 0.33333 | -1.8482 | 0 |
| 297 | 0.25475 | -2.6568 | 0 |
| 298 | 0.24748 | 0.81869 | 1 |
| 299 | 0.26088 | -2.4901 | 0 |
| 300 | 0.11111 | -3.6965 | 0 |
| 301 | 0.23981 | -3.7899 | 2 |
| 302 | 0.29971 | -2.7314 | 0 |
| 303 | 0.45781 | -1.5103 | 0 |
| 304 | 0.00015 | -14.786 | 0 |
| 305 | 0.24741 | -2.4901 | 0 |
| 306 | 0.12624 | -3.9035 | 0 |
| 307 | 0.33333 | -1.8482 | 0 |
| 308 | 0.00193 | 1.2334 | 2 |
| 311 | 0.22725 | -2.865 | 0 |
| 312 | 0.29902 | -2.0603 | 0 |
| 313 | 0.24741 | -2.4901 | 0 |
| 314 | 0.5046 | -1.3278 | 0 |
| **315** | **0.21062** | **-3.6929** | **3** |
| 318 | 0.32788 | -2.7169 | 1 |
| 319 | 0.15334 | -4.162 | 2 |
| 320 | 0.22712 | -3.9869 | 32 |
| 321 | 0.00686 | -10.165 | 1 |
| 322 | 0.03073 | -7.3382 | 28 |
| 323 | 0.22731 | -2.8642 | 0 |
| 325 | 0.24741 | -2.4901 | 0 |
| 327 | 0.00048 | -15.303 | 7 |
| 328 | 0.01195 | -8.5438 | 0 |
| 329 | 0.33333 | -1.8482 | 0 |
| 330 | 0.11111 | -3.6965 | 0 |
| 331 | 0.33333 | -1.8482 | 0 |
| 332 | 0.43366 | 1.63747 | 2 |
| 333 | 0.58216 | -0.837 | 3 |
| 334 | 0.29902 | -2.0603 | 0 |
| 336 | 0.24752 | 0.81872 | 4 |
| 337 | 0.57377 | -0.8527 | 3 |
| 338 | 0.50539 | 1.36257 | 1 |
| 341 | 0.06121 | -4.9802 | 0 |
| 342 | 0.33333 | -1.8482 | 0 |
| 343 | 0.08941 | -4.1207 | 0 |
| 344 | 0.24232 | -2.7541 | 0 |
| 345 | 0.00537 | -9.9603 | 0 |
| 346 | 0.01783 | -8.317 | 1 |
| 347 | 0.00015 | -14.786 | 0 |
| 349 | 0.19913 | -4.3902 | 7 |
| 350 | 0.33333 | -1.8482 | 0 |
| 351 | 0.0321 | -8.5923 | 7 |
| **353** | **0.43361** | **-1.6715** | **1** |
| 354 | 0.29902 | -2.0603 | 0 |
| 355 | 0.36127 | -1.8482 | 0 |
| 357 | 0.46228 | -1.5844 | 7 |
| 358 | 0.01235 | -7.3929 | 0 |
| 359 | 0.45386 | -1.5269 | 7 |
| **360** | **0.55404** | **1.85946** | **217** |
| 362 | 0.77995 | 0.9784 | 36 |
| 363 | 0.01701 | -10.645 | 35 |
| 364 | 0.03704 | -5.5447 | 0 |
| 365 | 0.26091 | 0.87901 | 1 |
| 366 | 0.46786 | -1.4329 | 1 |
| 368 | 0.29902 | -2.0603 | 0 |
| 371 | 0.06121 | -4.9802 | 0 |
| 372 | 0.2592 | -2.773 | 1 |
| 373 | 0.33333 | -1.8482 | 0 |
| 374 | 0.01481 | -7.3929 | 0 |
| 375 | 0.00318 | -12.57 | 0 |
| 376 | 0.00375 | -9.9603 | 0 |
| **377** | **0.33333** | **-1.8482** | **0** |
| 379 | 0.24741 | 0.81861 | 7 |
| 380 | 0.07326 | -4.9802 | 0 |
| 381 | 0.24741 | -2.4901 | 0 |
| 382 | 0.05423 | -5.7317 | 0 |
| 383 | 0.00137 | -11.089 | 0 |
| 384 | 0.1351 | -3.3685 | 0 |
| 385 | 0.0145 | -8.2104 | 0 |
| 386 | 0.33495 | -1.8393 | 0 |
| **387** | **0.24743** | **0.81867** | **1** |
| 389 | 0.01261 | -7.3777 | 0 |
| **390** | **0.01235** | **-7.3929** | **0** |
| 391 | 0.00093 | -12.45 | 0 |
| 392 | 0.01514 | -7.4702 | 0 |
| 393 | 0.43363 | -1.6714 | 1 |
| 395 | 0.01325 | -8.2534 | 0 |
| 397 | 0.12189 | -3.6965 | 0 |
| 398 | 0.00023 | -14.941 | 0 |
| 399 | 1.88E-06 | -22.179 | 0 |
| 400 | 0.01645 | -7.8731 | 0 |
| 402 | 0.51847 | -1.2809 | 0 |
| **403** | **0.04535** | **-7.3997** | **4** |
| 405 | 0.33333 | -1.8482 | 0 |
| 406 | 0.24741 | -2.4901 | 0 |
| 407 | 0.35538 | -1.8041 | 0 |
| 408 | 0.55546 | 1.8481 | 2 |
| 411 | 0.47862 | -1.3634 | 0 |
| 412 | 0.58857 | -0.7681 | 0 |
| 414 | 0.33329 | 0.92405 | 1 |
| 416 | 0.25964 | -2.6216 | 0 |
| 417 | 0.33333 | -1.8482 | 0 |
| 418 | 0.60593 | -0.6736 | 1 |
| 419 | 0.24741 | -2.4901 | 0 |
| 420 | 0.16811 | -4.2674 | 7 |
| 421 | 0.24741 | -2.4901 | 0 |
| 422 | 0.50567 | -1.3328 | 0 |
| 423 | 2.45E-06 | -24.624 | 0 |
| 424 | 0.33489 | -1.8397 | 0 |
| 425 | 0.33333 | -1.8482 | 0 |
| 426 | 0.33619 | -1.8482 | 0 |
| 428 | 0.11301 | -3.6965 | 0 |
| 429 | 0.03704 | -5.5447 | 0 |
| 430 | 0.25462 | -2.6556 | 0 |
| 431 | 5.08E-05 | -16.634 | 0 |
| 433 | 0.02621 | -8.4657 | 0 |
| 434 | 0.33333 | -1.8482 | 0 |
| 435 | 0.2346 | -2.8447 | 0 |
| 436 | 0.25206 | -2.8409 | 7 |
| 438 | 0.24743 | 0.81867 | 1 |
| 439 | 0.7257 | 1.20481 | 7 |
| 440 | 0.57416 | -0.8516 | 63 |
| 442 | 0.11111 | -3.6965 | 0 |
| 444 | 0.42936 | -3.1198 | 883 |

Most of the amino acid substitutions previously reported in PmrAB in isolates of *A. baumannii* that are resistant to colistin were found to be at non-polymorphic sites in our analysis. To date, four sites in *pmrA* and 26 sites in *pmrB* have been described as having non-synonymous mutations in the isolates resistant to colistin (**Supplementary Table S1**). Only one of the four resulting amino acid substitutions in PmrA and only three of the 26 in PmrB belong to polymorphic sites. In PmrA, the only substitution (S119T) at a polymorphic site, which has been shown to correlate with colistin resistance, occurs in the region that connects the domain of the signal receptor to the DNA binding domain (Figure 1**, top & middle panel**). In the case of PmrB, two of the three polymorphic sites in PmrB of colistin-resistant isolates have frequent substitutions A142V and P170Q; both sites fall into the intracellular domain of the protein. P360Q mutation occurs in the third polymorphic site, which lies in the ATPase domain of PmrB (Figure 2**, top & middle panel, Supplementary table S1**). Since the amino acid substitutions previously described in colistin resistant isolates of *A. baumannii* are sporadic observations, we understand that both PmrA and PmrB are constrained when accommodating non-synonymous SNPs. Therefore colistin resistance has not been an emerging phenomenon in the case of *A. baumannii*, unlike resistance to other antibiotics. We therefore examined the selection pressure at each amino acid site of the TCS, to see if there is negative selection acting on them, which could be the cause of the underlying mutation constraints.

### Both PmrA and PmrB are subject to negative selection pressure

To estimate the strength of natural selection pressure acting on each amino acid site in PmrA and PmrB, we performed codon based selection analysis. We observed significant negative selection (also known as ‘purifying selection’) all through both the protein sequences. In PmrA, 43 sites out of 224 are subjected to a strong negative selection pressure. As a result, these sites and the regions around them remain conserved (Figure 1, bottom panel). Functionally important sites, both in the signal receiver domain and in the DNA binding domain, exhibit a much more negative selection pressure than the rest of the sequence. While no signs of positive selection were found in PmrA, sites with high levels of polymorphism were found to be under neutral selection instead. Three of the four mutated sites of PmrA (M12K/M12I, P102H, S119T and L206P) that are reported in colistin-resistant isolates of *A. baumannii* belong to these highly polymorphic regions, which are under neutral selection (**Supplementary Table S1 &** Figure 1**, middle panel**).

Similarly, 52 out of 444 sites are under significant negative selection in PmrB (Table 3 & Figure 2, bottom panel). Sites under negative selection were well distributed throughout the protein sequence. As in the case of PmrA, these sites and regions around them are fairly conserved in PmrB. In particular, the extracellular region, which spans the functional sites of the HATPase_c, HisKA and HAMP domains, has far fewer mutations than the rest of the molecule, due to the presence of the majority of the sites under negative selection therein (Figure 2, top & middle panel). Notably, though most of these sites are under purifying selection, the functional domains managed to acquire amino acid substitutions (Figure 2, middle & bottom panel). This can explain the loss of fitness associated with colistin resistance in previous reports. As many as 23 out of the 26 amino acid substitutions in PmrB that occur in colistin-resistant isolates fall in functional domains (**Supplementary Table S1 &** Figure 2**, top panel**). However, these mutations occur mainly on sites that are under neutral selection (Figure 3A), each of them being delimited by the sites under negative selection in the functional domains (Figure 2**, top & bottom panels**). However, there exists a site at 153 of PmrB that is under significant positive selection (Table 3 **&** Figure 3B).

**Figure 3.**
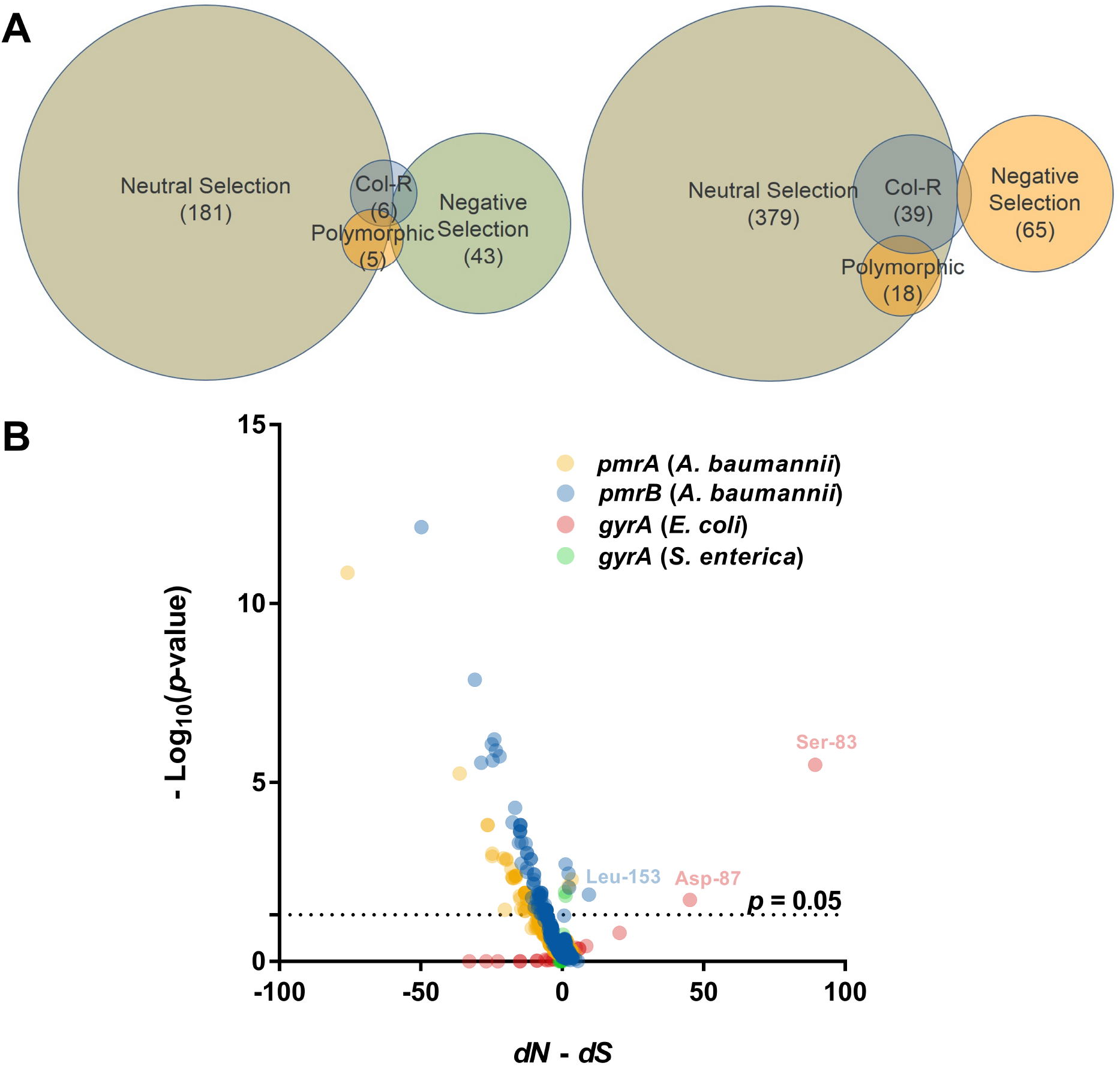
**A) Venn diagram depicting the amino acid substitutions resulting from non-synonymous mutations in** *pmrA* and *pmrB*, which are responsible for resistance to colistin in *A. baumannii* are under neutral selection, most of which are non-polymorphic sites. Numbers in each set indicate the number of sites falling under the corresponding category. **B)** Volcano plot of the *dN* – *dS* data for *pmrAB* and *gyrA* plotted against –Log_10_ of the *p*-value. The sites under adaptive selection in PmrB and GyrA have been indicated. This figure appears in colour in the online version of *JAC* and in black and white in the print version of *JAC*.

The SLAC method is equivalent to the site model of the equally well accepted program CodeML of the PAML package, but both employ different sets of procedures and algorithms (Yang, 2007). To examine the consistency of the obtained results, we performed the same selection analysis for PmrA and PmrB with CodeML. CodeML produced similar results to that we obtained with SLAC of HyPhy; it did not find any sign of significant positive selection in any of the sites of PmrA and PmrB (**Supplementary file S2**). This increases the reliability of the study and reinforces our results on the lack of adaptive selection on PmrAB.

Taken together, mutations in PmrAB TCS that are held responsible for colistin resistance in *A. baumannii* are neither adaptive nor polymorphic (Figure 3A). Rarher, the PmrAB TCS is under strong purifying selection to sustain colistin resistance in the long term. Therefore, it is plausible to believe that resistance to colistin, when occurred due to mutations in PmrAB, is not disseminated owing to the underlying purifying selection that causes the loss of fitness.

### Selection analysis of gyrA of E. coli and S. enterica was able to detect signatures of adaptive evolution that validates our approach

To validate this approach of selection analysis and increase reliability of the results, we employed the same methodology to detect positively selected sites in well-known antibiotic targets that cause antibiotic resistance. We considered DNA gyrase A (*gyrA*) to be the most appropriate for this purpose, as there is abundant of literature that share common findings about mutations in this gene in multiple organisms that lead to quinolone resistance. Mutations at Ser-83 and Asp-87 in GyrA are the most frequently reported sites to be responsible for quinolone resistance in Enteric bacteria (Domínguez et al., 2002; Everett et al., 1996; Lindgren et al., 2003; Vila et al., 2001, 1994). Therefore, our results of selection analysis can be qualified as valid if these two sites are detected under the same selection analysis framework applied to *pmrAB*. Interestingly, selection analysis of *gyrA* successfully detected these two sites (Ser-83 and Asp-87) to be under adaptive evolution in both *E. coli* as well as *S. enterica* (Figure 3B **& Supplementary file S2**). This not only substantiated the reliability of the methodology, but also validated our results obtained for *pmrAB*. In addition, this also provided evidences that adaptive evolution had an important role to play behind the emergence of quinolone resistance in *E. coli* and *S. enterica*, while it has little role to play in colistin resistance in *A. baumannii*.

## Discussion

The emergence and spread of multidrug-resistant ESKAPE group of pathogens including *A. baumannii* have prompted many health professionals to assume that we are on the verge of a post-antibiotic era. Colistin and to some extent, tigecycline appear to be the only treatment options for treating infections caused by multidrug-resistant *A. baumannii*. In addition, the notable efficacy of colistin against tigecycline-resistant *A. baumannii* advocates colistin as the last option against the organism. However, reports of colistin resistance in *A. baumannii* have raised unprecedented concerns, as it is thought that resistance to colistin may become widespread in the future if not used judiciously. Contrary to this belief, we have shown in this work that resistance to colistin may not become a general phenomenon and these concerns are therefore unwarranted. Similar opinions were also put forwarded by López-Rojas *et al.* based on the observed reduced fitness and reduced virulence of experimentally induced colistin-resistant *A. baumannii* isolates (López-Rojas et al., 2011). However, they also cautioned that, though the synonymous mutations are initially not adaptive, compensatory mutations may attenuate the reduced fitness. In contrast, using selection analysis, our investigation provided more direct evidence that mutations that cause colistin resistance are neither adaptive nor polymorphic and therefore, may not take over the global population.

In *A. baumannii*, resistance to colistin occurs in two mechanisms: enzymatic modification of lipid A moieties of the bacterial cell wall, either by the addition of phosphoethanolamine or galactosamine, or hyperacylation, and complete loss of cell wall lipo-polysaccharide (LPS) due to inactivation of lipid A biosynthesis (Beceiro et al., 2011; Boll et al., 2015; Henry et al., 2011; Moffatt et al., 2011, 2010; Pelletier et al., 2013). Colistin resistance by both mechanisms is the result of deletions and/or substitutions in the PmrAB two-component system (Adams et al., 2009; Arroyo et al., 2011; Beceiro et al., 2011). Resistance to colistin due to mutations in PmrAB is the most common mode of colistin resistance.

PmrAB mutation analysis, in conjunction with domain analysis suggests that the signal receptor domain and the DNA binding domain, as well as the functional sites therein are ideally conserved in PmrA. This indicates that PmrA transduces a very specific signal, input from PmrB and in turn, regulates a limited and non-degenerate set of genes. On the other hand, the transmembrane domains as well as the intracellular domain of PmrB are highly variable. Other domains such as the kinase and ATP-binding domain are comparatively conserved and contain more sites with purifying selection. Since there is only one intracellular domain delimited by transmembrane α-helices, this domain must be recruiting the response regulator(s). Since this domain is variable, we can state that it recruits more than one response regulators and/or transcription factors, resulting in cross-talks between parallel signal transduction pathways, one of which is the cognate PmrA-mediated signalling. More detailed investigations at the molecular level are needed to corroborate this hypothesis.

In this study, we collected 3113 draft genomes from the RefSeq database to evaluate the degree of natural selection acting on each site of PmrAB two-component system. Initially, the study considered *pmrAB* sequences only related to colistin resistant strains. This comprise of 44 *pmrA* and 39 *pmrB* sequences that were collected from the whole genomes of colistin resistant strains as well as from the full length coding sequences associated with colistin resistance that appear in the literature. Selection analysis with this dataset showed just one site with purifying selection and no adaptive selection in PmrA. Similarly, it showed neither positive nor negative selection in PmrB (**Supplementary file S2**). However, when all the *pmrAB* sequences from the 3113 genomes were considered, it was found that the PmrAB TCS is under strong purifying selection. This dramatic change in the results can be attributed to the homogeneity of the initial sequence data that contains sequences associated with only colistin resistant strains. In cases where the sequences belong to a homogenous population exposed to similar environmental conditions, most of the mutations are transient mutations, which underestimates any real purifying as well as adaptive selection (Kryazhimskiy & Plotkin, 2008). A similar scenario exists when the strains undergo convergent and parallel evolution resulting from colistin exposure (Crandall et al., 1999). The larger dataset allowed the inclusion of genomes reported worldwide over a wide range of time scale that ensures a heterogeneous set of samples/populations for analysis.

To evaluate long term emergence of colistin resistance, we need to examine the scenario post-exposure to colistin. In addition, if any site is under adaptive selection, selection analysis should identify signatures of adaptive selection from a sufficiently large dataset, even in the absence of colistin exposure. We could then qualify these mutations are stable enough to sustain long term resistance to colistin. Most of the amino acid substitutions in PmrAB reported to date are either from longitudinal isolates of patients or induced in laboratory conditions. Reports of clinical isolates bearing colistin resistance are scant. In addition, the colistin exposure to *A. baumannii* lasts as long as the treatment of the patient, and the resistance (if arises) reverts back to susceptible phenotype on withdrawal of colistin. (Adams et al., 2009; Cheah et al., 2016; Snitkin et al., 2013). Therefore, it is plausible to believe that mutations that cause colistin resistance are transient mutations owing to the short exposure to the antibiotic as clinicians prefer to give short term colistin therapy due to its association with nephrotoxicity. Further, with no evidence of compensatory mutations linked to colistin resistance identified till date, which strengthens our supposition that colistin resistant mutations may never get fixed in the global population of *A. baumannii* (Lesho et al., 2013; López-Rojas et al., 2011; Rolain et al., 2011). This observation may not apply to the populations in micro-environments, such as a laboratory cultures.

Apparently, resistance to colistin is achieved by a cost of fitness. Studies in the past have shown reduced fitness in colistin resistant isolates, both in terms of growth rate under laboratory conditions and reduced virulence (Beceiro et al., 2014; López-Rojas et al., 2011; López-Rojas et al., 2013). In addition, to compensate for the loss/modification of the LPS, the transcriptome and the corresponding proteome are largely shifted towards expression of genes responsible for membrane integrity and biogenesis (Boll et al., 2016; Cheah et al., 2016; Henry et al., 2011). A recent work confirms that surface lipoproteins are overexpressed and localized profusely on the cell surface, which could compensate for the loss of membrane integrity. These studies highlight the excessive cost of colistin resistance, and therefore, mutations in PmrAB may be deleterious and subject to strong purifying selection.

A significant number of sites in both PmrA and PmrB are under purifying selection. In addition, we also identified mutation hotspots, sites that are exceptionally polymorphic. Interestingly, sites with mutations that cause colistin resistance are not polymorphic sites. Rather, colistin resistance mutations are largely under neutral selection. In addition, a few of colistin resistance mutations are also under purifying selection (Figure 3A & B). We suggest that these conditions maintain the incidence of colistin resistance under check. Further, loss of fitness due to colistin resistance resulting from the combined effects of neutral colistin resistance mutations and the purifying selection acting on the overall protein sequences ensure that the colistin resistance remains uncommon. The reasons for this loss of fitness include physiological costs in terms of slow growth rate, loss of membrane integrity and its compensation, and reduced virulence. In addition to this, because of the high selection pressure exerted by the host’s immune system and the presence of drugs, the short exposure of the bacterium to colistin in the host, it is highly unlikely that random genetic drift can fix these mutations. These observations explain us why reports of colistin-resistant isolates of *A. baumannii* have been sporadic and they might remain so. In the near future, *A. baumannii* may not be able to emerge as a colistin-resistant organism, unless other mechanisms such as plasmid-mediated (*e. g.*, *mcr-*1) resistance emerge.

The other mechanism of development of Colistin resistance is due to mutations in *lpxACD* of *A. baumannii*. However, unlike mutations in *pmrAB*, where virtually all mutations that cause Colistin resistance are due to substitutions, mutations reported in *lpxACD* are due to large indels, non-sense mutations and insertion sequence-mediated gene disruptions that cause complete loss of cell wall lipopolysaccharide (Carretero-Ledesma et al., 2018; Girardello et al., 2017; Moffatt et al., 2010). There is no direct method to analyse signatures of selection pressure from such mutations. However, it has already been reported through growth curve and virulence assays that fitness costs associated with *lpx* mutations are even higher than those associated with *pmr* mutations (Beceiro et al., 2014) and therefore, *lpxACD* mutants are unsustainable in the long run.

One site at 153 of PmrB appears to be under adaptive selection. This site has *dN* – *dS* value of 9.48851 (*p* = 0.013). Surprisingly, this site has never been reported to be mutated in colistin resistant strains of *A. baumannii*. This site belongs to one of the transmembrane domains of PmrB. The fact that it has so far never caused colistin resistance indicates that the positive selection gain may not overcome the cost of colistin resistance. Since the site belongs to a transmembrane domain, it is also possible that the mutations that occur at this site might serve an advantage in terms of structural stability, rather than having a functional advantage. Further, studies employing site directed mutagenesis experiments could throw additional information about this site.

The study attempted to assert the reliability of the methodology by applying it to *gyrA* on the basis of whether it can identify sites that are expected to be under adaptive selection in multiple organisms. The computational method of selection analysis has precisely detected the sites under adaptive evolution in GyrA, an observation which is supported by a wealth of previous literature about quinolone resistance in bacteria. In case of *E. coli*, resistance to quinolones requires multiple mutations in different genes including *gyrA* to achieve a clinically relevant level. The requirement for multiple mutations suggests that there is strong selection for resistance mutations invariably associated with unusually high mutation rates occurring due to antibiotic exposure. Quinolones are known to interact with the DNA-GyrA complex, rather than with the enzyme alone. One study correlated that *gyrA* mutations lead to reduction of drug binding to the protein-DNA complex to 60-fold less than that of wild-type GyrA (Willmott & Maxwell, 1993). In this respect, identification of the sites that are most pronounced to be associated with quinolone resistance by adaptive selection analysis provides sufficient grounds to emphasize the power of the methodology to detect adaptive evolution in protein coding sequences.

## Conclusion

The anticipation and management of natural selection is an important aspect of the fight against multidrug resistance (Lesho et al., 2013). This was the central theme of this work in the context of the degree of natural selection acting on PmrAB TCS, which has been implicated in resistance to colistin. We showed that in case of resistance to colistin, natural selection plays only a minor role, if at all, and therefore, continuing use of colistin could still be a safe option. Therefore, the concerns raised by healthcare professionals regarding rapid emergence of colistin resistance and about continuing the usage of colistin for the treatment against *A. baumannii* infections are unwarranted.

## Acknowledgements

We duly thank and acknowledge the remote access to supercomputing provided by **Bioinformatics Resources and Applications Facility (BRAF), C-DAC, Pune** for the computationally intensive analyses.

## Funding

This work was supported by University Grants Commission, Government of India [Grant Ref. No. F. 30-70/2016 (SA-II)].

## Transparency declarations

None to declare.

## Author contributions

**SP**: Literature search, Figures, Data collection, Data analysis, Bench works, Writing of the article; **RS, AR**: Data collection, Bench works, Data analysis; **AT**: Critical revision of the article, administrative, technical, or logistic support; **KPA**: Conception and design, analysis and interpretation, writing of the article, provision of materials, patients, and resources; **PK**: Supervision of the entire work, Conceptual design of the experiment, analysis and interpretation, writing of the article, provision of materials, patients, resources and funding.

## Nucleotide sequence accession numbers

Nucleotide sequences of *pmrAB* from *Acinetobacter baumannii* strains PKAB19 and PKAB15 have been deposited in GenBank under accession numbers MH925082 and MH925083, respectively.

## Notes

#### Summary of Updates

The title was changed from "Darwinian selection analysis of the two-component system PmrAB indicates there could be lingering delay in emergence of colistin resistance in Acinetobacter baumannii" to "Paucity of adaptive selection in PmrAB two-component system may resist emergence of colistin resistance in Acinetobacter baumannii"

