## Supplementary methods for "Paucity of adaptive selection in PmrAB two-component system may resist emergence of colistin resistance in *Acinetobacter baumannii*"

***Antibiotic susceptibility testing***

Minimum inhibitory concentration (MIC) for colistin were performed according to European Committee on Antimicrobial Susceptibility Testing guidelines (EUCAST, 2017). E-test (Hi-media, Mumbai) was used for colistin MIC determination. The reference strain *A. baumannii* ATCC 19606 and quality control strain *Escherichia coli* ATCC 25922 were included in the assays as reference controls.

***Mutational analysis of pmrA***

Mutations in *pmrAB* have been known to be associated with colistin resistance. We amplified *pmrA* genes from two isolates namely PKAB15 and PKAB19 from our collection (Table 1), which showed higher MIC values using the *in house* designed primers 5'-AAG CCA ACA AAC TAA ACA AAA-3' and 5'-GCT TGC TCA ACA GGT GGA AC-3' and the amplicons generated were custom sequenced (Macrogen Inc., Seoul, South Korea). The origin, molecular characters and other clinical details of the isolates were given in **Table 1**.

***Literature search for mutations reported in colistin resistant isolates***

This study reviewed previous literatures that have identified non-redundant and non-synonymous mutations in *pmrAB* in colistin resistant *A. baumannii* isolates (up to December 2018) (**Supplementary Table S1**). Reports of *pmrAB* point mutations without any mention of corresponding colistin resistance phenotypes were excluded from this study. Amino acid substitutions reported in colistin resistant phenotypes that appear in colistin sensitive strains were also excluded. The mutations found in the literature were mapped onto the amino acid sequence and analysed in the context of natural selection at the respective sites of the mutations.

***Selection analysis of pmrAB***

To estimate the degree of the selection pressure acting on each amino acid site in PmrA and PmrB, we performed selection analysis and expressed it in terms of the difference between the rates of non-synonymous and synonymous mutations (*dN* – *dS*). Given a codon-aligned multiple nucleotide sequence alignment corresponding to a protein sequence, the selection analysis counts the number of non-synonymous mutations (*dN*) at each amino acid position and compares it with the number of synonymous mutations (*dS*) at that site, tested on a phylogenetic tree. The extent of selection pressure at a codon can be estimated by comparing the rate of occurrence of synonymous mutations to that of non-synonymous mutations. The extent of selection pressure at a codon can be estimated by comparing the rate of occurrence of synonymous mutations (*dS*) to that of non-synonymous mutations (*dN*). For a site under neutral selection, synonymous and non-synonymous mutations will occur at similar rate and therefore, *dN* ≈ *dS*. However, if a site is subjected to high selective pressure, the site is under purifying selection, which allows only synonymous mutations, in which case *dN* < *dS*. On the other hand, a site with adaptive selection will gain more non-synonymous substitutions than synonymous mutations, resulting in *dN* > *dS*.

In total, 3113 draft genome sequences of *A. baumannii* (as available up to December 2018) were retrieved from National Centre for Biotechnology Information RefSeq database. Genomes of strains of *A. baumannii* isolated during time-span of more than 30 years (1984 - 2018) were included in the analysis and such a long-term of isolation period should be sufficient enough to answer a query of evolution in *A. baumannii.* The dosage used for those resistant strains during	treatment was 80–160 mg every 8 h for >60 kg bodyweights that may reach serum concentration of up to a maximum of 5mg/L, while in *in-vitro* experimental studies it ranged from 1 to >256 μg/μl. Thus, it includes resistances that have arisen both during patients’ treatment as well as by laboratory induction.

Coding sequences for *pmrA* and *pmrB* were extracted from a standalone BLAST+ database of the downloaded genome sequences by ‘tBLASTn-Fast’ search algorithm of BLAST+ v2.8.1. An E-value cut-off of 1e-10 and ‘max_target_seqs’ value (maximum target sequences to keep) of 3113 were kept for the tBLASTn search. Hits with <90% identity were filtered out from tabular output before extracting the coding sequences. Sequences with indels and truncated / partial sequences were removed manually prior to alignment. Only complete coding sequences free from indels were included in the study. The resulting coding sequences were utilized to perform codon based multiple sequence alignment by MEGA7 software with the MUSCLE algorithm (Edgar, 2004; Kumar et al., 2016). For this purpose, the coding sequences were translated into protein sequences, followed by multiple protein sequence alignment and reverse-translation of the alignments into nucleotide sequences. As a result, the alignment was based on triplet codons instead of individual nucleotides, which would allow identification of synonymous and non-synonymous mutations.

The codon-based multiple sequence alignment files were introduced into the Datamonkey server for selection analysis. The Datamonkey server is the web interface for the HyPhy package for the analysis of molecular evolution (Delport et al., 2010; Sergei L. Kosakovsky Pond & Frost, 2005; Pond & Muse, 2005). Single-Likelihood Ancestor Counting (SLAC) method was chosen for the estimation of synonymous and non-synonymous mutation rates at each site. SLAC was preferred to other options based on its simplicity and speed, as well as its performance on closely related sequences (Pond & Frost, 2005). The neighbor joining tree based on HKY85 nucleotide substitution model, which was the best fit model, have been chosen for the estimation of branch lengths and substitution rates for SLAC (Hasegawa et al., 1985). Model comparison is an implicit automated procedure of the web-server, which as described in Pond et al. (Pond & Frost, 2005). A *p*-value of <0.05 from a two-tailed extended binomial distribution was used as a cut-off for the significance of positive selection against a neutral null model. Positive selection was detected in terms of the difference between non-synonymous and synonymous mutations, which was expressed as *dN* – *dS* value, normalized with the total length of the phylogenetic tree as measured in terms of the number of expected substations per nucleotide per site. A site with amino acid mutations in >1% (≈30) genomes was considered as polymorphic. Results of codon-wise mutation and selection analysis and the corresponding *p*-values are provided in **Supplementary file S2.** The codon-aligned multiple sequence alignment files of all *pmrA* and *pmrB* coding sequences are available in **Supplementary file S3.**

The SLAC method is equivalent to the site model of the CodeML program of the PAML package (Yang, 2007). Therefore, we performed an adaptive selection analysis under the site model (Model M2a), against a nearly-neutral evolution null model (Model M1a) in CodeML (PAML v4.9) (Jeffares et al., 2015; Yang, 2007). Briefly, the file was parsed into RaxML for the construction of phylogenetic tree with ‘ML + Rapid bootstrap’ option and under general time reversible-gamma-invariant model (GTRGAMMAI) with 1000 bootstrap repeats. The bootstrap and branch length values were kept enabled (‘BS brl’ enabled) (Stamatakis, 2014). *pmrA* of *A. junii* strain 65 and *pmrB* of *A. nosocomialis* strain M2 were used as outgroups for the phylogenetic analysis of *pmrA* and *pmrB* of *A. baumannii*, respectively. The tree files and the sequence files were used as inputs to CodeML. The phylogenetic trees, sequence files, CodeML control files and the result files have been provided in **Supplementary file S3**.

To demonstrate that our approach of selection analysis is consistent and reliable, we intended to apply the same methodology to a gene of other bacteria that is thought to be under adaptive evolution in the context of antibiotic resistance. We chose the gene coding for DNA gyrase subunit A (*gyrA*) of *Escherichia coli* and *Salmonella enterica* , the product of which is the target for quinolone class of antibiotics, for this purpose (Hershberg, 2017; Katz & Hershberg, 2013). We applied within species codon-based selection analysis as described above on *gyrA* sequences of 533 genomes of *E. coli* and 553 genomes of *S. enterica* randomly downloaded from the Refseq genome database. The results can be found in **Supplementary File S2**, whereas raw alignments can be found in **Supplementary file S4**.

**Nucleotide sequence accession numbers**

Nucleotide sequences of *pmrAB* from *Acinetobacter baumannii* strains PKAB19 and PKAB15 have been deposited in GenBank under accession numbers MH925082 and MH925083, respectively.
