## Supplementary Table S1 for "Paucity of adaptive selection in PmrAB two-component system may resist emergence of colistin resistance in *Acinetobacter baumannii*"

Mutations that are associated with colistin resistance, collected from the scientific literature.

| **Gene** | **Mutation** | **Reference** |
| --- | --- | --- |
| *pmrB* | M145K, P233S, L87F, S14L, F387Y, S403F, A227V, N353Y | (Beceiro et al., 2011) |
|  | P233S | (Dahdouh et al., 2017) |
|  | P233S, R263C, M145I, T13A | (Snitkin et al., 2013) |
|  | D64V, L208F, P170Q, P170L, R263P, R263C, A226V, P233S, T235I, N256I,A80V, G315D, Q277H, P360Q, P377L | (Arroyo et al., 2011) |
|  | A262P, A227V, T13N, P233S, P233T | (Adams et al., 2009) |
|  | R134C, G315S, Y194S | (Thi Khanh Nhu et al., 2016) |
|  | A227V | (Cheah et al., 2016) |
|  | A142V | (Haeili et al., 2018) |
|  | T234I, H264Y, L208F, I339M, V390E, S17R, P360Q | (Napier et al., 2013) |
|  | S17R, T232I, R263L, Y116H | (Napier et al., 2013) |
|  | H263R, V444I, I235T, G390V | (Lee et al., 2016) |
|  | P233S | (Durante-Mangoni et al., 2015) |
|  | S17R, T235I | (Wand et al., 2015) |
|  | P233S, P170L | (Pournaras et al., 2014) |
|  | G272D | (Mu et al., 2016) |
|  | A227V, P233S | (Kim et al., 2014) |
|  | I121F, A183T, A184V P190S, T192I, Q228P | (Park et al., 2011) |
| *pmrA* | L206P | (Snitkin et al., 2013) |
|  | M12I, S119T | (Arroyo et al., 2011) |
|  | P102H | (Adams et al., 2009) |
|  | M12K | López-Rojas et al., 2013; Dortet et al., 2018 |
|  | E8D | (Napier et al., 2013; Rolain et al., 2013) |
|  | M12R | (Lee et al., 2016) |
|  | G54E | (Oikonomou et al., 2015) |
